# GOcats: A tool for categorizing Gene Ontology into subgraphs of user-defined concepts

**DOI:** 10.1101/306936

**Authors:** Eugene W. Hinderer, Hunter N.B. Moseley

**Author notes:** **Software and full results available at:** http://software.cesb.uky.edu https://figshare.com/s/134d06901a7a6f331934 (Scripts used to generate results) https://github.com/MoseleyBioinformaticsLab/GOcats (software version 1.1.4) https://pypi.python.org/pypi/GOcats (software version 1.1.4) http://gocats.readthedocs.io/en/latest/index.html (documentation) https://figshare.com/s/21defede0cd3865742d4 (Figures and Supplementary Data).

## Abstract

Gene Ontology is used extensively in scientific knowledgebases and repositories to organize the wealth of available biological information. However, interpreting annotations derived from differential gene lists is difficult without manually sorting into higher-order categories. To address these issues, we present GOcats, a novel tool that organizes the Gene Ontology (GO) into subgraphs representing user-defined concepts, while ensuring that all appropriate relations are congruent with respect to scoping semantics. We tested GOcats performance using subcellular location categories to mine annotations from GO-utilizing knowledgebases and evaluating their accuracy against immunohistochemistry datasets in the Human Protein Atlas (HPA).

In comparison to mappings generated from UniProt’s controlled vocabulary and from GO slims via OWLTools’ Map2Slim, GOcats outperforms these methods without reliance on a human-curated set of GO terms. By identifying and properly defining relations with respect to semantic scope, GOcats can use traditionally problematic relations without encountering erroneous term mapping. We applied GOcats in the comparison of HPA-sourced knowledgebase annotations to experimentally-derived annotations provided by HPA directly. During the comparison, GOcats improved correspondence between the annotation sources by adjusting semantic granularity. Utilized in this way, GOcats can perform an accurate knowledgebase-level evaluation of curated HPA-based annotations.

## Introduction

### Gene Ontology (GO)

The Gene Ontology (GO) [1] is the most common biological controlled vocabulary (CV) used to represent information and knowledge distilled from almost every type of biological and biomedical research data generated today, from classic wet-bench experiments to high-throughput analytical platforms, especially omics technologies. Each CV term in GO is assigned a unique alphanumeric code, which is used to annotate genes and gene products in many other databases, including UniProt [2] and Ensembl [3]. GO is divided into three sub-ontologies: Cellular Component, Molecular Function, and Biological Process. A graph can represent each sub-ontology, where individual GO terms are nodes connected by directional edges (i.e. relation). For example, the term “lobed nucleus” (GO:0098537) is connected by a directional is_a relation edge to the term “nucleus” (GO:0005634). In this graph context, the is_a relation defines the term “nucleus” as a parent of the term “lobed nucleus”. There are eleven types of relations used in the core version of GO; however, is_a is the most ubiquitous. The three GO sub-ontologies are “is_a disjoint” meaning that there are no is_a relations connecting any node among the three sub-ontologies.

There are also three versions of the GO database: *go-basic* which is filtered to exclude relations that span across sub-ontologies and to include only relations that point toward the top of the full ontology; *go* or *go-core* contains additional relations, that may span sub-ontologies and which point both toward and away from the top of the ontology; and *go-plus* contains cross-references to entries in external databases and ontologies.

### Growth and Evolution of Biological Controlled Vocabularies

GO and other CVs like the Unified Medical Language System [4,5] saw an explosion in development in the mid-1990s and early 2000s, coinciding with the increase in high-throughput experimentation and “big data” projects like the Human Genome Project. Their intended purpose was, and still is, to standardize the functional descriptions of biological entities so that these functions can be referenced via annotations across large databases unambiguously, consistently, and with increased automation. However, as large-scale and high-throughput investigations continue to advance, ontologies are also evolving as they are used in ways that extend beyond their initial purpose as annotation reference utilities. Ontology annotations are utilized alongside automated pipelines that analyze protein-protein interaction networks and form predictions of unknown protein function based on these networks [6,7], for gene function enrichment analyses, and are now being leveraged for the creation of predictive disease models in the scope of systems biochemistry [8].

### Difficulty in representing biological concepts derived from omics-level research

Differential abundance analyses for a range of omics-level technologies, especially transcriptomics technologies can yield large lists of differential genes, gene-products, or gene variants. Many different GO annotation terms may be associated with these differential gene lists, making it difficult to interpret without manually sorting into appropriate descriptive categories [9]. It is similarly non-trivial to give a broad overview of a gene set or make queries for genes with annotations of a biological concept. For example, a recent effort to create a protein-protein interaction network analysis database resorted to manually building a hierarchical localization tree from GO cellular compartment terms due to the “incongruity in the resolution of localization data” in various source databases and the fact that no published method existed at that time for the automated organization of such terms [6]. If subgraphs of GO could be programmatically extracted to represent such concepts, a category-defining general term could be easily associated with all its ontological child terms. These category-defining terms would enable a more robust and easily interpretable organization of genes and gene products for the investigation of specific biological and disease processes and facilitate the development of complex biological models such as bio-macromolecular interaction and metabolic networks.

Meanwhile, high-throughput transcriptomic and proteomic characterization efforts like those carried out by the Human Protein Atlas (HPA) now provide sophisticated pipelines for resolving expression profiles at organ, tissue, cellular and subcellular levels by integrating quantitative transcriptomics with microarray-based immunohistochemistry [10]. Such efforts are creating a huge amount of omics-level experimental data that is cross-validated and distilled into systems-level annotations linking genes, proteins, biochemical pathways, and disease phenotypes across our knowledgebases. However, annotations provided by such efforts may vary in terms of granularity, annotation sets used, or ontologies used. Therefore, (semi-)automated and unbiased methods for categorizing semantically-similar and biologically-related annotations are needed for integrating information from heterogeneous sources—even if the annotation terms themselves are standardized—to facilitate effective downstream systems-level analyses and integrated network-based modeling.

### Term categorization approaches

Issues of term organization and term filtering have led to the development of GO slims—manually trimmed versions of the gene ontology containing only generalized terms [11] which represent concepts within GO, as well as other software, like Categorizer [9], which can organize the rest of GO into representative categories using semantic similarity measurements between GO terms. GO slims may be used in conjunction with mapping tools, such as OWLTools’ Map2Slim (M2S), or GOATools [12,13], to map fine-grained annotations within Gene Annotation Files (GAFs) to the appropriate generalized term(s) within the GO slim or within a list of GO terms of interest. While web-based tools such as QuickGO exist to help compile lists of GO terms [14], using Map2Slim either relies completely on the structure of existing GO slims or requires input or selection of individual GO identifiers for added customization, and necessitates the use of other tools for mapping. UniProt has also developed a manually-created mapping of GO to a hierarchy of biologically-relevant concepts [15]. However, it is smaller and less maintained than GO slims, and is intended for use only within UniProt’s native data structure.

In addition to utilizing the inherent hierarchical organization of GO to categorize terms, other metrics may be used for categorization. For instance, semantic similarity can be combined along with the GO structure to calculate a statistical value indicating whether a term should belong to a predefined group or category of [9,16–19]. One rationale for this type of approach is that the topological distance between two terms in the ontology graph is not necessarily proportional to the semantic closeness in meaning between those terms, and semantic similarity reconciles potential inconsistencies between semantic closeness and graph distance. Additionally, some nodes have multiple parents, where one parent is more closely related to the child than the others [9]. Semantic similarity can help determine which parent is semantically more closely related to the term in question. While these issues are valid, we maintain that in the context of aggregating fine-grained terms into general categories, these considerations are not necessary. First, fluctuations in semantic distances between individual terms are not an issue once terms are binned into categories: all binned terms will be reduced to a single step away from the category-defining node. Second, the problem of choosing the most appropriate parent term for a GO term only causes problems when selecting a representative node for a category; however, since most paths eventually converge onto a common ancestor, any significantly diverging paths would have its meaning captured by rooting multiple categories to a single term, cleanly sidestepping the issue.

### Maintenance of ontologies

Despite maintenance and standard policies for adding terms, ontological organization is still subject to human error and disagreement, necessitating quality assurance and revising, especially as ontologies evolve or merge. A recent review of current methods for biomedical ontology mapping highlights the importance in developing semi-automatic methods to aid in ontology evolution efforts and reiterates the aforementioned concept of semantic correspondence in terms of scoping between terms [20]. Methods incorporating such correspondences have been published elsewhere, but these deal with issues of ontology evolution and merging, and not with categorizing terms into user-defined subsets [21,22]. Ontology merging also continues to be an active area of development for integrating functional, locational, and phenotypic information. To aid in this endeavor, another recent review points out that it is crucial to integrate phenotypic information across various levels of organismal complexity, from the cellular level to the organ system level [8]. Thus, organizing location-relevant ontology terms into discrete categories is an important step toward this end.

### GO Categorization Suite (GOcats)

For the reasons indicated above, we have developed a tool called the GO Categorization Suite (GOcats), which serves to streamline the process of slicing the ontology into custom, biologically-meaningful subgraphs representing concepts derivable from GO. Unlike previously developed tools, GOcats uses a list of user-defined keywords and/or GO terms that describe a broad category-representative term from GO, along with the structure of GO and augmented relation properties to generate a subgraph of child terms and a mapping of these child terms to their respective category-defining term that is automatically identified based on the user’s keyword list, or to the GO term that is explicitly specified. Furthermore, these tools allow the user to choose between the strict axiomatic interpretation or a looser semantic scoping interpretation of part-whole (mereological) relation edges within GO. Specifically, we consider scoping relations to be comprised of is_a, part_of, and has_part, and mereological relations to be comprised of part_of and has_part.

Here, we demonstrate the utility of GOcats and the effectiveness of evaluating mereological relations with respect to semantic scope by categorizing the GO Cellular Component ontology into broad-level concepts representing cellular components. We used the concept-centric subgraphs produced by GOcats to create a mapping of fine-grained terms to their chosen concept-representative term. Using these mappings, we categorized knowledgebase-derived gene annotations and compared this automated categorization to publicly available datasets of manually-categorized gene annotations assigned by researchers at the HPA following immunohistochemistry experiments.

## Design and Implementation

GOcats is designed to use the go-core version of the GO database, which contains other relations that connect the separate ontologies and may point away from the top of the ontology. GOcats can either exclude non-scoping relations or invert has_part directionality into a part_of_some interpretation, maintaining the acyclicity of the graph. Therefore, GOcats can represent *go-core* as a directed acyclic graph (DAG).

GOcats is a Python package written in major version 3 of the Python program language [23] and available on GitHub and the Python Package Index (PyPI). It uses a Visitor design pattern implementation [24] to parse the *go-core* Ontology database file [4]. Searching with user-specified sets of keywords for each category, GOcats extracts subgraphs of the GO DAG (sub-DAGs) and identifies a representative node for each category in question and whose child nodes are detailed features of the components. Figure 1 illustrates this approach in more detail. Pseudocode and additional descriptions for the algorithm are provided in Supplementary Data 1.

**Figure 1.**
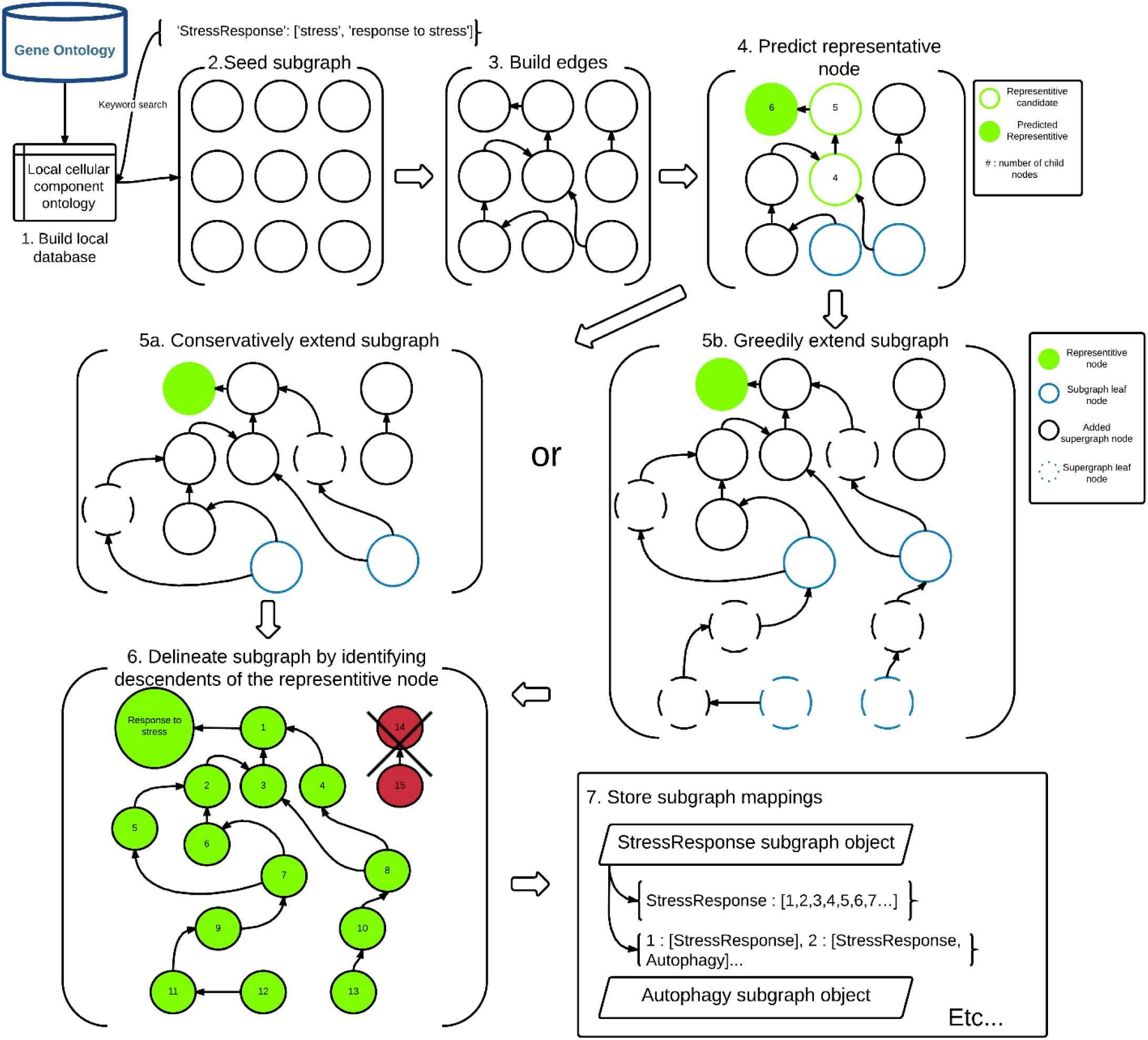
Flowchart of the GOcats’ subgraph creation method.

To overcome the previously mentioned issues regarding scoping ambiguity among mereological relations, we assigned properties indicating which term was broader in scope and which term was narrower in scope to each edge object created from each of the scope-relevant relations in GO. For example, in the node pair connected by a part_of or is_a edge, node 1 is narrower in scope than node 2. Conversely, node 1 is broader in scope than node 2 when connected by a has_part edge. This edge is therefore reinterpreted by GOcats as part_of_some. We have published additional explanations for this re-interpretation elsewhere and demonstrate improvement in annotation enrichment statistics when this re-interpretation is used [Link to sister manuscript].

While the default scoping relations in GOcats are is_a, part_of, and has_part, the user has the option to define the scoping relation set. For instance, one can create go-basic-like subgraphs from a go-core version ontology by limiting to only those relations contained in go-basic. For convenience, we have added a command line option, “go-basic-scoping,” which allows only nodes with is_a and part_of relations to be extracted from the graph.

For mapping purposes, Python dictionaries are created which map GO terms to their corresponding category or categories. For inter-sub-DAG analysis, another Python dictionary is created which maps each category to a list of all its graph members. By default, fine-grained terms do not map to category root-nodes that define a sub-DAG that is a superset of a category with a root-node nearer to the term. For example, a member of the “nucleolus” sub-DAG would map only to “nucleolus,” and not to both “nucleolus” and “nucleus”. However, the user has the option to override this functionality if needed. Mapping supersets is a requirement for visualizing concept membership in graph representations using tools like Cytoscape (Figure 2).

**Figure 2.**
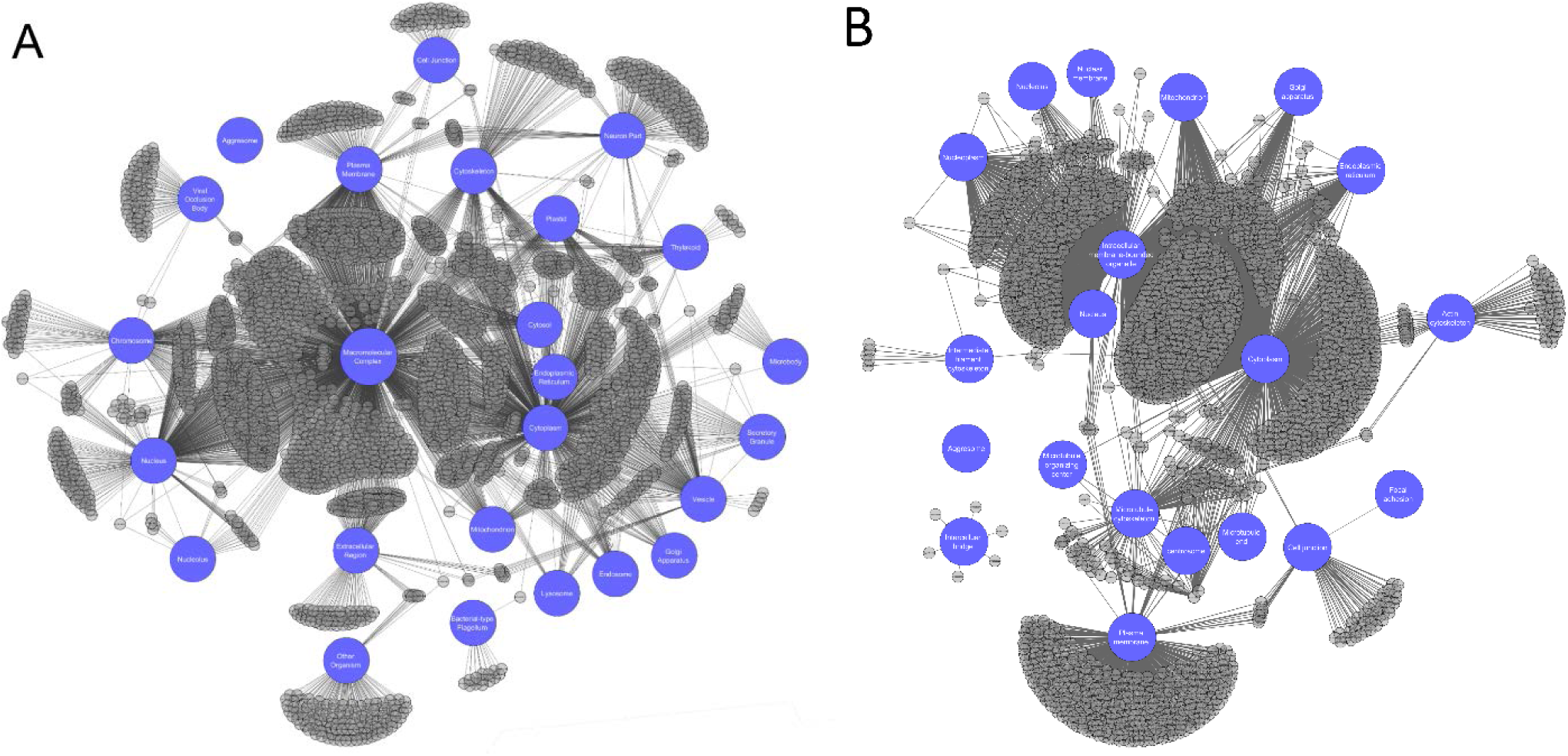
Extracted GO subcellular localization subgraphs form a network with expected connectivity patterns. *Blue nodes indicate category-representative nodes and grey nodes represent fine-grained GO terms that are part of their respective subgraphs. Edges connect fine-grained terms to their GOcats-assigned representative node(s). Images were created using Cytoscape 3.0* [29]. *A) Network of 25 categories whose subgraphs account for 89% of the GO cellular component sub-ontology* B) Network of 20 categories used in the Human Protein Atlas subcellular localization immunohistochemistry raw data.

## Results

### GOcats compactly organizes GO subcellular localization terms into user-specified categories

We evaluated the automatic extraction and categorization of 25 subcellular locations, using GOcats’ “comprehensive” method of subgraph extension (Figure 1, Supplementary Data 1). Of these, 22 contained a designated GO term root-node that exactly matched the concept intended at the creation of the keyword list (Table 1). These subgraphs account for approximately 89% of GO’s Cellular Component sub-ontology. While keyword querying of GO provided an initial seeding of the growing subgraph, Table 1 highlights the necessity of re-analyzing the GO graph to find terms missed by the keyword search, to remove terms erroneously added by the keyword search, and to add appropriate subgraph terms not captured by the keyword search. For example, the “cytoplasm” subgraph grew from its initial seeding of 296 nodes to 1197 nodes after extension. Conversely, 136 nodes were seeded by keyword for the “bacterial” subgraph, but only 16 were truly rooted to the representative node.

**Table 1:**
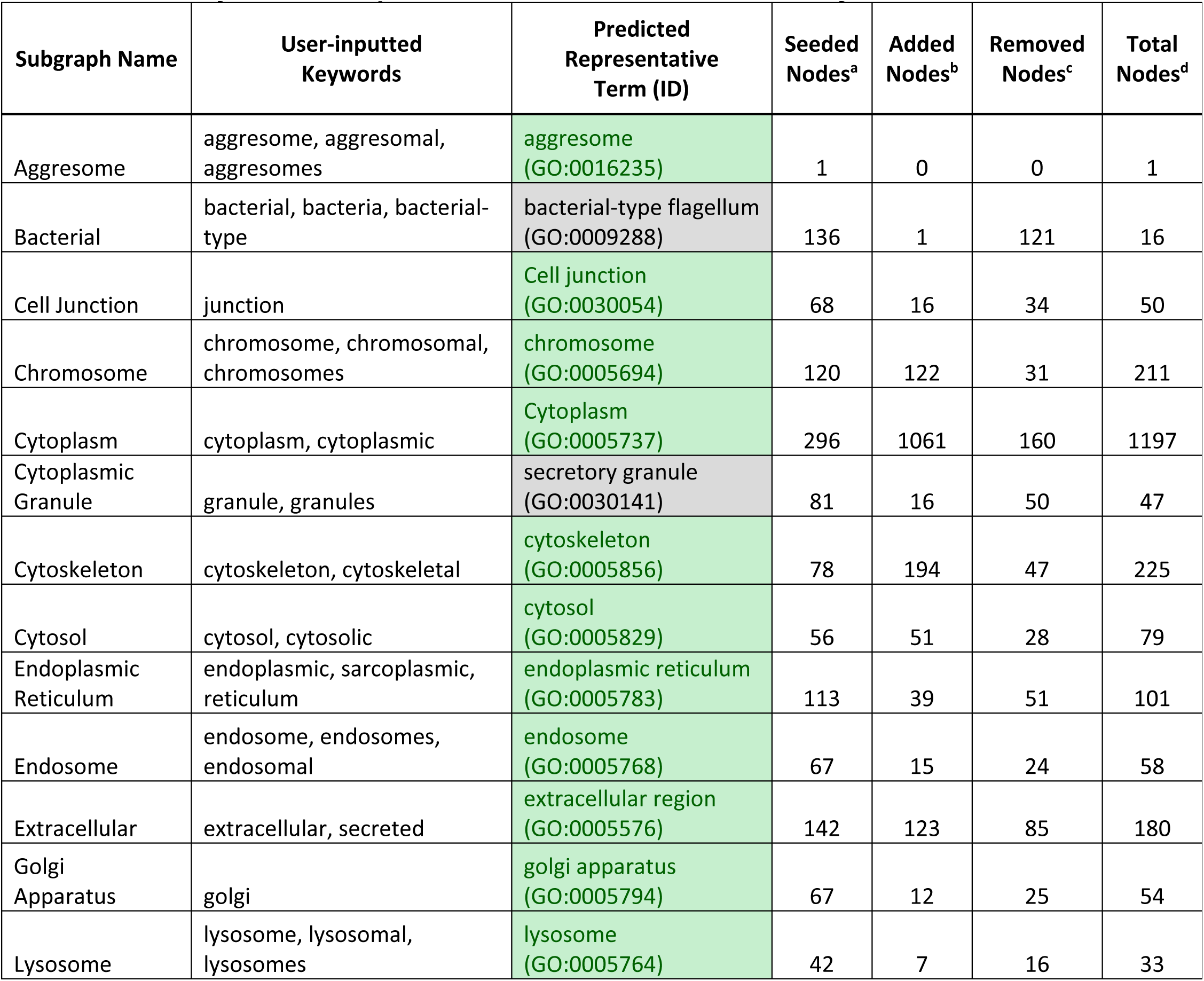
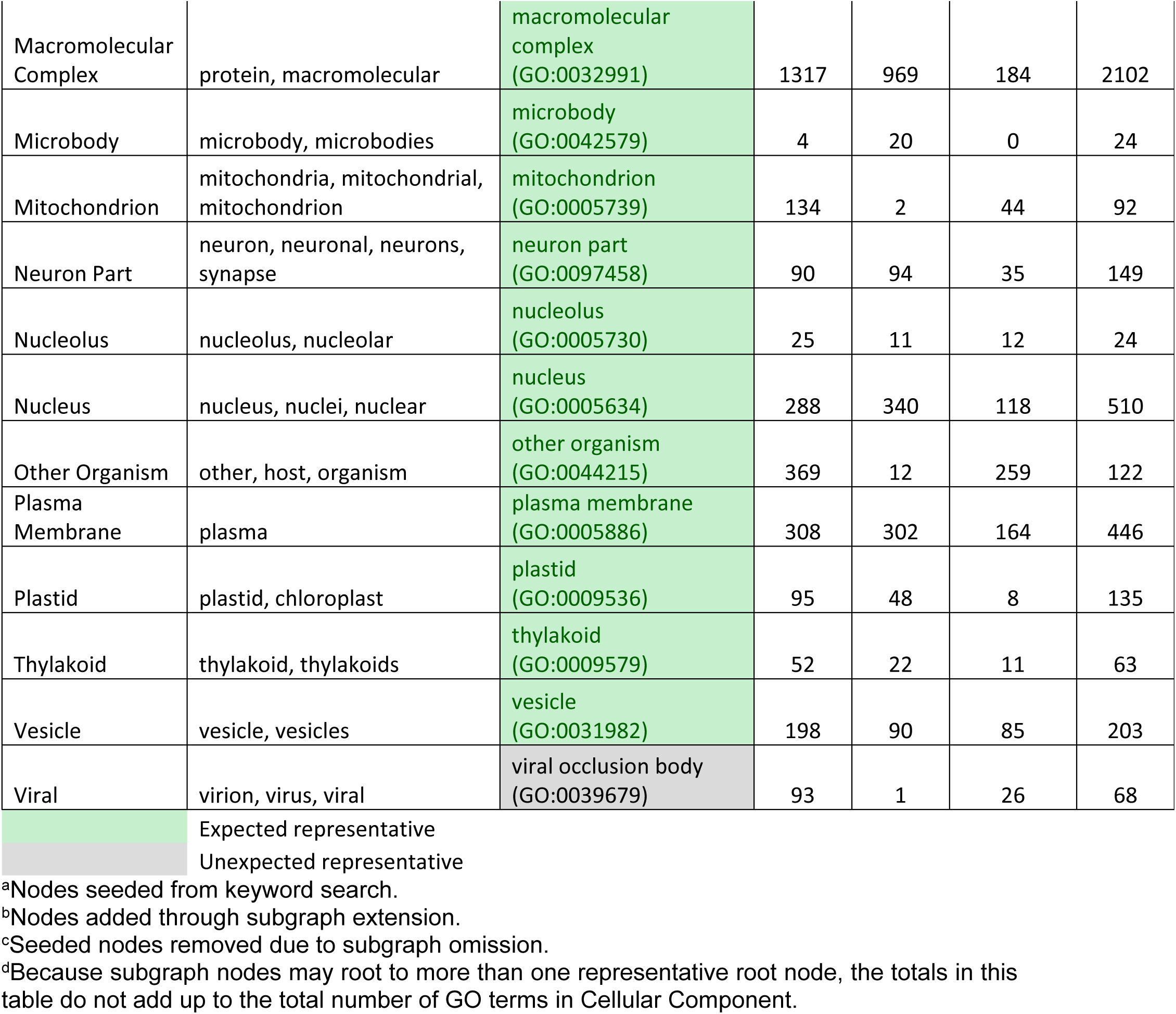
Summary of 25 example subcellular locations extracted by GOcats

| Subgraph Name | User-inputted Keywords | Predicted Representative Term (ID) | Seeded Nodes<sup>a</sup> | Added Nodes<sup>b</sup> | Removed Nodes<sup>c</sup> | Total Nodes<sup>d</sup> |
| --- | --- | --- | --- | --- | --- | --- |
| Aggresome | aggresome, aggresomal, aggresomes | aggresome (GO:0016235) | 1 | 0 | 0 | 1 |
| Bacterial | bacterial, bacteria, bacterial-type | bacterial-type flagellum (GO:0009288) | 136 | 1 | 121 | 16 |
| Cell Junction | junction | Cell junction (GO:0030054) | 68 | 16 | 34 | 50 |
| Chromosome | chromosome, chromosomal, chromosomes | chromosome (GO:0005694) | 120 | 122 | 31 | 211 |
| Cytoplasm | cytoplasm, cytoplasmic | Cytoplasm (GO:0005737) | 296 | 1061 | 160 | 1197 |
| Cytoplasmic Granule | granule, granules | secretory granule (GO:0030141) | 81 | 16 | 50 | 47 |
| Cytoskeleton | cytoskeleton, cytoskeletal | cytoskeleton (GO:0005856) | 78 | 194 | 47 | 225 |
| Cytosol | cytosol, cytosolic | cytosol (GO:0005829) | 56 | 51 | 28 | 79 |
| Endoplasmic Reticulum | endoplasmic, sarcoplasmic, reticulum | endoplasmic reticulum (GO:0005783) | 113 | 39 | 51 | 101 |
| Endosome | endosome, endosomes, endosomal | endosome (GO:0005768) | 67 | 15 | 24 | 58 |
| Extracellular | extracellular, secreted | extracellular region (GO:0005576) | 142 | 123 | 85 | 180 |
| Golgi Apparatus | golgi | golgi apparatus (GO:0005794) | 67 | 12 | 25 | 54 |
| Lysosome | lysosome, lysosomal, lysosomes | lysosome (GO:0005764) | 42 | 7 | 16 | 33 |
| Macromolecular Complex | protein, macromolecular | macromolecular complex<br>(GO:0032991) | 1317 | 969 | 184 | 2102 |
| Microbody | microbody, microbodies | microbody<br>(GO:0042579) | 4 | 20 | 0 | 24 |
| Mitochondrion | mitochondria, mitochondrial, mitochondrion | mitochondrion<br>(GO:0005739) | 134 | 2 | 44 | 92 |
| Neuron Part | neuron, neuronal, neurons, synapse | neuron part<br>(GO:0097458) | 90 | 94 | 35 | 149 |
| Nucleolus | nucleolus, nucleolar | nucleolus<br>(GO:0005730) | 25 | 11 | 12 | 24 |
| Nucleus | nucleus, nuclei, nuclear | nucleus<br>(GO:0005634) | 288 | 340 | 118 | 510 |
| Other Organism | other, host, organism | other organism<br>(GO:0044215) | 369 | 12 | 259 | 122 |
| Plasma Membrane | plasma | plasma membrane<br>(GO:0005886) | 308 | 302 | 164 | 446 |
| Plastid | plastid, chloroplast | plastid<br>(GO:0009536) | 95 | 48 | 8 | 135 |
| Thylakoid | thylakoid, thylakoids | thylakoid<br>(GO:0009579) | 52 | 22 | 11 | 63 |
| Vesicle | vesicle, vesicles | vesicle<br>(GO:0031982) | 198 | 90 | 85 | 203 |
| Viral | virion, virus, viral | viral occlusion body<br>(GO:0039679) | 93 | 1 | 26 | 68 |

|  |  |
| --- | --- |
|  | Expected representative |
|  | Unexpected representative |

<sup>a</sup>Nodes seeded from keyword search.
<sup>b</sup>Nodes added through subgraph extension.
<sup>c</sup>Seeded nodes removed due to subgraph omission.
<sup>d</sup>Because subgraph nodes may root to more than one representative root node, the totals in this table do not add up to the total number of GO terms in Cellular Component.

Of note, 2102 of the 3877 terms in Cellular Component could be rooted to a single concept: “macromolecular complex.” Despite cytosol being defined as “the part of the cytoplasm that does not contain organelles, but which does contain other particulate matter, such as protein complexes,” less than half of the terms rooted to macromolecular complex also rooted to cytosol or cytoplasm. Surprisingly, approximately 25% of the terms rooted to macromolecular complex are rooted to this category alone and the remaining are rooted only to macromolecular complex and another compartment that was extracted.

The visualization of the subgraph contents confirmed the uniqueness of the macromolecular complex category and showed the relative sizes of groups of GO terms shared between two or more categories (Figure 2A). Overall, the patterns of connectedness in this network make sense biologically, within the constraints of GO’s internal organization (Supplementary Data 2).

### GOcats robustly categorizes GO terms into category subgraphs with high similarity to existing GO-utilizing categorization methods while including information gleaned from has_part edges

To assess the accuracy of GOcats’ category subgraph contents, we evaluated the similarity of these subgraph contents to subgraphs of the manually-curated UniProt subcellular localization CV [2,15] (see Supplementary Data 3 and 4). In comparing the overlap of terms between UniProt’s CV and corresponding category-representative nodes produced by GOcats, most GOcats-derived subgraphs are large supersets of UniProt subgraphs. Supplementary Data 5 shows that 12 of the GOcats-derived compartments had identical root nodes in UniProt’s considerably smaller controlled vocabulary. Of these, 6 contained 100% inclusion and were approximately 20 times larger on average. The others contained between 56.2% and 84.6% inclusion. Some discrepancies in the organizational patterns between UniProt and GO may account for the lower inclusion. One major discrepancy is UniProt’s organization of the plasma membrane and other cellular envelopes.

We also performed comparisons with subgraphs created by M2S (Supplementary Data 6 and 7). This method is more comparable to GOcats, because it directly utilizes the GO graph structure. In comparing the category subgraphs created by GOcats and M2S (Supplementary Data 8), the mappings for most categories are in very close agreement, as evidenced by both high inclusion and Jaccard indices in Supplementary Data 9 and further highlighted in Figures 3A, 3B and Supplementary Data 10 A-V [25]. However, in some categories, M2S and GOcats disagree as illustrated in Figure 3C and Supplementary Data 10E. The most striking example of this is in the plasma membrane category, where M2S’s subgraph contained over 300 terms that were not mapped by GOcats. We manually examined theses discrepancies in the plasma membrane category and noted that many of the terms uniquely mapped by M2S did not appear to be properly rooted to “plasma membrane” (Supplementary Data 11).

**Figure 3.**
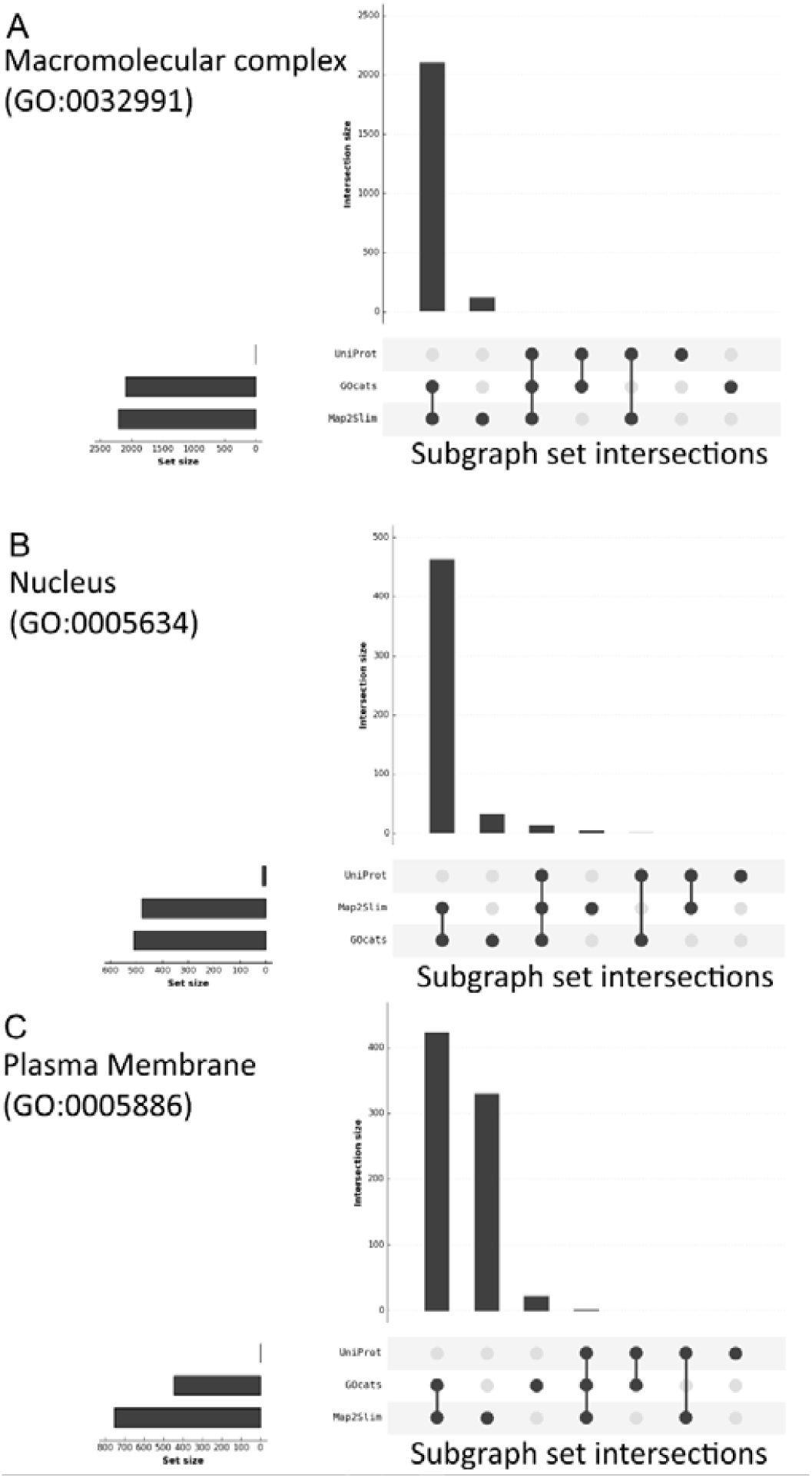
Visualizing the degree of overlap between the category subgraphs created by GOcats, Map2Slim, and the UniProt CV. Plots were created using the R package: UpSetR [25] and represent the subgraphs created from mapping fine-grained terms in GO to the indicated general category using the indicated mapping method for: A) Macromolecular Complex; B) Nucleus; C) Plasma Membrane. Plots for all categories can be found in Supplementary Data 10A-Y.

M2S mapped terms such as “nuclear envelope,” “endomembrane system,” “cell projection cytoplasm”, and “synaptic vesicle, resting pool” to the plasma membrane category, while such questionable associations were not made using GOcats. Even though most terms included by M2S but excluded by GOcats exist beyond the scope of or are largely unrelated to the concept of “plasma membrane,” a few terms in the set did seem appropriate, such as “intrinsic component of external side of cell outer membrane.” However, of these examples, no logical semantic path could be traced between the term and “plasma membrane” in GO, indicating that these associations are not present in the ontology itself. **These differences in mapping are due to our reevaluation of the has_part edges with respect to scope.** As shown in Supplementary Data 9 the categories with the greatest agreement between the two methods were those with no instances of has_part relations, which is the only relation in Cellular Component that is natively incongruent with respect to scope. However, there is no apparent correlation between the frequency of this relation and the extent of disagreement.

### Custom-tailoring of GO slim-like categories with GOcats allows for robust knowledgebase gene annotation mining

The ability to query knowledgebases for genes and gene products related to a set of general concepts-of-interest is an important method for biologists and bioinformaticians alike. Using the set of GO terms annotated in the HPA’s immunohistochemistry localization raw data as “concepts” (Table 2), we derived mappings to annotation categories generated from GOcats, M2S, and UniProt’s CV based on UniProt-and Ensembl-sourced annotations from the European Molecular Biology Laboratories-European Bioinformatics Institute (EMBL-EBI) QuickGO knowledgebase resource [14] (Supplementary Data 12). These annotation-category mappings were also visualized using Cytoscape (Figure 2B). Next, we evaluated how these derived annotation categories matched raw HPA data GO annotations (overall analysis illustrated in Supplementary Data 13). GOcats slightly outperformed M2S and significantly outperformed UniProt’s CV in the ability to query and extract genes and gene products from the knowledgebase that exactly matched the annotations provided by the HPA (Figure 4A). Similar relative results are seen for partially matched knowledgebase annotations. Genes in the “partial agreement,” “partial agreement is superset,” or “no agreement” groups may have annotations from other sources that place the gene in a location not tested by the HPA immunohistochemistry experiments or may be due to non-HPA annotations being at a higher semantic scoping than what the HPA provided. Also, novel localization provided by the HPA could explain genes in the “partial agreement” and “no agreement” groups. Furthermore, GOcats performed the categorization of HPA’s subcellular locations dataset in 5.971 seconds when filtered to the cellular localization sub-ontology and 9.248 seconds when unfiltered, while M2S performed its mapping on the same data in 13.393 seconds. Although comparable, GOcats should offer appreciable computational improvement on significantly larger datasets. This is rather surprising since GOcats is implemented in Python [23], an interpreted language, versus M2S which is implemented in Java and compiled to Java byte code.

**Table 2:** Summary of 20 subcellular locations used in the HPA raw experimental data extracted by GOcats

| Subgraph Name | User-inputted Keywords | Predicted Representative Term (ID) | Seeded Nodes<sup>a</sup> | Added Nodes<sup>b</sup> | Removed Nodes<sup>c</sup> | Total Nodes<sup>d</sup> |
| --- | --- | --- | --- | --- | --- | --- |
| Actin cytoskeleton | actin cytoskeleton | actin cytoskeleton (GO:0015629) | 117 | 22 | 77 | 62 |
| Aggresome | aggresome, aggresomal, aggresomes | aggresome (GO:0016235) | 1 | 0 | 0 | 1 |
| Cell Junction | junction | cell junction (GO:0030054) | 68 | 16 | 34 | 50 |
| Centrosome | centrosome | centrosome (GO:0005813) | 10 | 2 | 5 | 7 |
| Cytoplasm | cytoplasm, cytoplasmic | cytoplasm (GO:0005737) | 296 | 1061 | 160 | 1197 |
| Endoplasmic Reticulum | endoplasmic, sarcoplasmic, reticulum | endoplasmic reticulum (GO:0005783) | 113 | 39 | 51 | 101 |
| Focal adhesion | focal adhesion | focal adhesion (GO:0005925) | 29 | 0 | 28 | 1 |
| Golgi Apparatus | golgi | golgi apparatus (GO:0005794) | 67 | 12 | 25 | 54 |
| Intercellular bridge | intercellular bridge | intercellular bridge (GO:0045171) | 24 | 2 | 19 | 7 |
| Intermediate filament cytoskeleton | intermediate filament cytoskeleton | intermediate filament cytoskeleton (GO:0045111) | 126 | 0 | 118 | 8 |
| Intracellular membrane-bounded organelle (vesicle)<sup>e</sup> | intracellular membrane-bounded organelle | Intracellular membrane-bounded organelle (GO:0043231) | 229 | 1116 | 118 | 1227 |
| Microtubule cytoskeleton | microtubule cytoskeleton | microtubule cytoskeleton (GO:0015630) | 112 | 55 | 68 | 109 |
| Microtubule end | microtubule end | microtubule end (GO:1990752) | 138 | 0 | 133 | 5 |
| Microtubule organizing center | microtubule organizing center | microtubule organizing center (GO:0005815) | 110 | 34 | 95 | 49 |
| Mitochondrion | mitochondria, mitochondrial, mitochondrion | mitochondrion (GO:0005739) | 134 | 2 | 44 | 92 |
| Nuclear membrane | nuclear membrane | nuclear membrane (GO:0031965) | 1151 | 0 | 1139 | 12 |
| Nucleolus | nucleolus, nucleolar | nucleolus (GO:0005730) | 25 | 11 | 12 | 24 |
| Nucleoplasm | nucleoplasm | nucleoplasm (GO:0005654) | 10 | 125 | 4 | 131 |
| Nucleus | nucleus, nuclei, nuclear | nucleus (GO:0005634) | 288 | 340 | 118 | 510 |
| Plasma Membrane | plasma | plasma membrane (GO:0005886) | 308 | 302 | 164 | 446 |

Expected representative
Unexpected representative
<sup>a</sup>Nodes seeded from keyword search.
<sup>b</sup>Nodes added through subgraph extension.
<sup>c</sup>Seeded nodes removed due to subgraph omission.
<sup>d</sup>Because subgraph nodes may root to more than one representative root node, the totals in this table do not add up to the total number of GO terms in Cellular Component.
<sup>e</sup>HPA conservatively annotates "vesicles" as intracellular membrane-bounded organelle.

**Figure 4.**
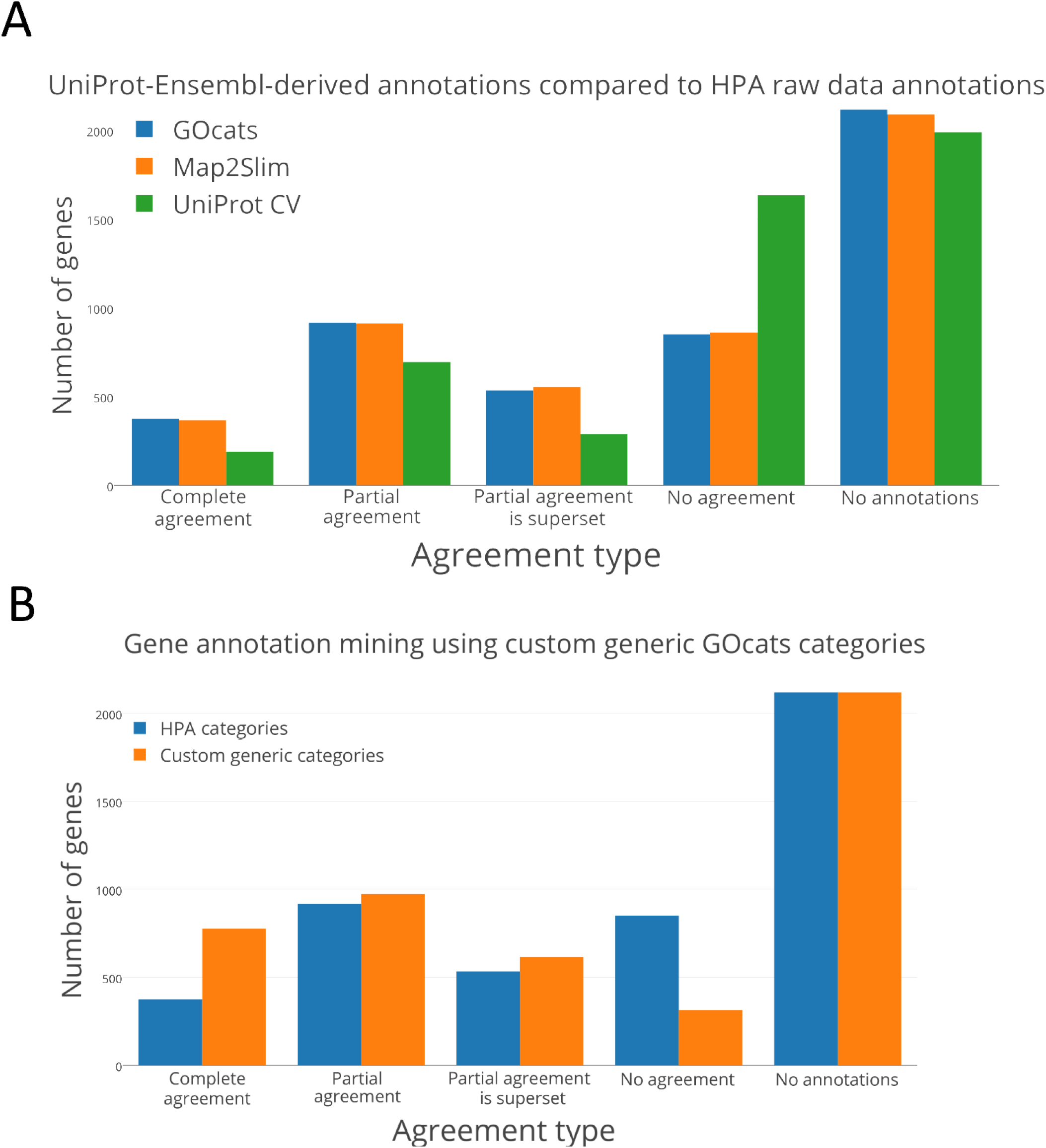
Comparison of UniProt-Ensembl knowledgebase annotation data mining extraction performance by GOcats, Map2Slim, and UniProt CV. “Complete agreement” refers to genes where all subcellular locations derived from the knowledgebase and the HPA dataset matched, “partial agreement” refers to genes with at least one matching subcellular location, “partial agreement is superset” refers to genes where knowledgebase subcellular locations are a superset of the HPA dataset (these are mutually exclusive to the “partial agreement” category), “no agreement” refers to genes with no subcellular locations in common, and “no annotations” refers to genes in the experimental dataset that were not found in the knowledgebase. The more-generic categories used in panel B can be found in Table 3. (continued below) A) Number of genes of the given agreement type when comparing mapped gene product annotations assigned by UniProt and Ensembl in the EMBL-EBI knowledgebase to those taken from The Human Protein Atlas’ raw data. Knowledgebase annotations were mapped by GOcats, Map2Slim, and the UniProt CV to the set of GO annotations used by the HPA in their experimental data. B) Shift in agreement following GOcats’ mapping of the same knowledgebase gene annotations and the set of annotations used in the raw experimental data using a more-generic set of location terms meant to rectify potential discrepancies in annotation granularity.

One key feature of GOcats is the ability to easily customize category subgraphs of interest. To improve agreement and rectify potential differences in term granularity, we used GOcats to organize HPA’s raw data annotation along with the knowledgebase data into slightly more generic categories (Table 3). In doing so, GOcats can query over twice as many knowledgebase-derived gene annotations with complete agreement with the more-generic HPA annotations, while also increasing the number of genes in the categories of “partial” and “partial agreement is superset” agreement types and decreasing the number of genes in the “no agreement” category (Figure 4B).

**Table 3:** Generic location categories used to resolve potential scoping inconsistencies in HPA raw data

| HPA annotation category | GOcats-customized general HPA category |
| --- | --- |
| Actin cytoskeleton | Cytoskeleton |
| Centrosome |  |
| Intermediate filament cytoskeleton |  |
| Microtubule cytoskeleton |  |
| Microtubule end |  |
| Microtubule organizing center |  |
| Aggresome | Aggresome |
| Cell junction | Cell junction |
| Cytoplasm | Cytoplasm |
| Endoplasmic reticulum | Endoplasmic reticulum |
| Focal adhesion | Focal adhesion |
| Golgi apparatus | Golgi apparatus |
| Intercellular bridge | intercellular bridge |
| intracellular membrane-bounded organelle | intracellular membrane-bounded organelle |
| Mitochondrion | Mitochondrion |
| Nucleus | Nucleus |
| Nucleoplasm |  |
| Nuclear membrane |  |
| Nucleolus | Nucleolus |
| Plasma membrane | Plasma membrane |

We then compared the methods’ mapping of knowledgebase gene annotations derived from HPA to the HPA experimental dataset to demonstrate how researchers could use the GOcats suite to evaluate how well their own experimental data is represented in public knowledgebases. Because the set of gene annotations used in the HPA experimental dataset and in the HPA-derived knowledgebase annotations are identical, no term mapping occurred during the agreement evaluation and so the assignment agreement was identical between GOcats and M2S. As expected, the complete agreement category was high, although there was a surprising number of partial agreement and even some genes that had no annotations in agreement (Figure 5). We next broke down which locations were involved in each agreement type and noted that the “nucleus,” “nucleolus,” and “nucleoplasm” had the highest disagreement relative to their sizes, but that disagreements were present across nearly all categories (Supplementary Data 14).

**Figure 5.**
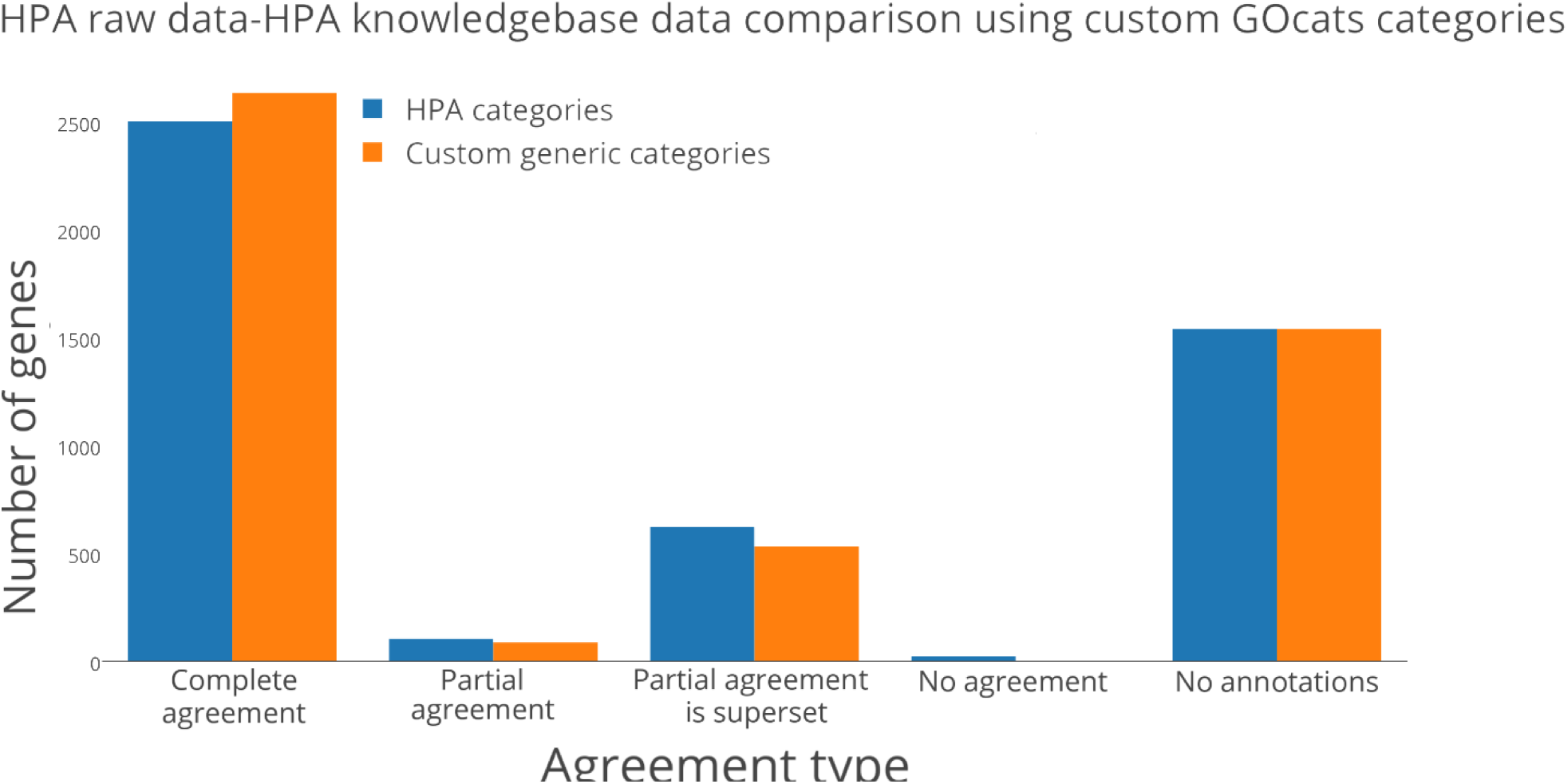
Comparison of HPA knowledgebase derived annotations to HPA experimental data. Number of genes in the given agreement type when comparing gene product annotations assigned by HPA in the EMBL-EBI knowledgebase to those in The Human Protein Atlas’ raw experimental data. “Complete agreement” refers to genes where all subcellular locations derived from the knowledgebase and the HPA dataset matched, “partial agreement” refers to genes with at least one matching subcellular location, “partial agreement is superset” refers to genes where knowledgebase subcellular locations are a superset of the HPA dataset (these are mutually exclusive to the “partial agreement” category), “no agreement” refers to genes with no subcellular locations in common, and “no annotations” refers to genes in the experimental dataset that were not found in the knowledgebase. The more-generic categories used in panel B can be found in Table 3.

Both M2S and GOcats avoid superset category term mapping; neither map a category-representative GO term to another category-representative GO term if one supersedes another (although GOcats has the option to enable this functionality). Therefore, discrepancies in annotation should not arise by term mapping methods. Nevertheless, we hypothesized that some granularity-level discrepancies exist between the HPA experimental raw data and the HPA-assigned gene annotations in the knowledgebase. We performed the same custom category generic mapping as we did for the previous test and discovered that some disagreements were indeed accounted for by granularity-level discrepancies, as seen in the decrease in “partial” and “no agreement” categories and increase in “complete” agreement category following generic mapping (Figure 5, blue bars). For example, 26S proteasome non-ATPase regulatory subunit 3 (PSMD3) was annotated to the nucleus (GO:0005634) and cytoplasm (GO:0005737) in the experimental data but was annotated to the nucleoplasm (GO:0005654) and cytoplasm in the knowledgebase. By matching the common ancestor mapping term “nucleus”, GOcats can group the two annotations in the same category. In total, 132 terms were a result of semantic scoping discrepancies. Worth noting is the fact that categories could be grouped to common categories to further improve agreement, for example “nucleolus” within “nucleus.”

Interestingly, among the remaining disagreeing assignments were some with fundamentally different annotations. Many of these are cases in which either the experimental data, or knowledgebase data have one or more additional locations distinct from the other. For example, NADH dehydrogenase [ubiquinone] 1 beta subcomplex subunit 6 (NDUB6) was localized only to the mitochondria (GO:0005739) in the experimental data yet has annotations to the mitochondria and the nucleoplasm (GO:0005654) in the knowledgebase. Why such discrepancies exist between experimental data and the knowledgebase is not immediately clear.

We were also surprised by the high number of genes with “supportive” annotations in the HPA raw data that were not found in the EMBL-EBI knowledgebase when filtered to those annotated by HPA. As Figure 5 shows, roughly one-third of the annotations from the raw data were missing altogether from the knowledgebase; the gene was not present in the knowledgebase whatsoever. This was surprising because “supportive” was the highest confidence score for subcellular localization annotation.

## Discussion

In this study, we: i) demonstrated the increase in retrievable ontological information content via reevaluating mereological relations to make them congruent with respect to semantic scope, ii) applied our new method GOcats toward the categorization and utilization of the GO Cellular Component sub-ontology, and iii) evaluated the ability of GOcats and other mapping tools to relate HPA experimental to HPA knowledgebase GO Cellular Component annotation sources.

Our results indicate that, when compared to UniProt’s CV, GOcats’ mapping was able to assign gene annotations from the UniProt and Ensembl database to subcellular locations with greater accuracy when compared to a raw dataset of gene localization annotations. It is important to note that UniProt’s CV was not intended to be used as a method to categorize gene annotations. Nevertheless, it is itself a DAG with a structure comparable to GO and we analyzed the graph and mapped fine-grained terms to general terms using the same techniques used for GO. Moreover, GOcats comparison to M2S demonstrates similar mapping performance between the two methods, but with GOcats providing important improvements in mapping, computational speed, ease of use, and flexibility of use. Using GOcats, the user can create custom, GO slim-like filters to map fine-grained gene annotations from GAFs to general subcellular compartments without needing to hand-select a set GO terms for categorization. Moreover, users can use GOcats to quickly customize the level of semantic specificity for annotation categories. We have used this functionality to automatically map gene annotations from Ensembl and UniProt-GOA knowledgebases and compared these localization assignments to manually-assigned localizations taken from high-throughput immunohistochemistry experiments performed by the HPA [13]. We show that GOcats allows a robust organization of Cellular Component into user-specified categories, while providing more automation than current methods.

Furthermore, we demonstrate GOcats’ ability to query and organize gene-specific annotations from knowledgebases into experimentally-verified general subcellular locations with safer scoping utilization over other methods that require specific versions of GO. Finally, we demonstrate GOcats’ utility for evaluating annotation assignment consistency between raw experimental data and knowledgebase data, highlighting this software’s promise for knowledgebase curation and quality control.

Other methods used to summarize or categorize GO terms into biologically relevant concepts: i) rely heavily on static and manually-maintained GO slims, ii) require the user to create GO slims by hand-selecting GO terms and creating a GO slim from scratch, or iii) require the user to perform post-analysis categorization of GO terms, typically enriched GO terms from gene enrichment studies. Caveats of such methods include burdening the user with the tasks of finding and editing, or even creating the appropriate GO slims to suit their research, limiting categorization to only those concepts that are explicitly defined in GO, and in the case of post-enrichment categorization tools, limiting the statistical power of the analysis by not automatically binning gene annotations into categories prior to enrichment. Until GOcats, there has been no resource developed which categorizes GO terms into subsets representing concepts without user-specification of individual GO terms or the use of GO slims, which operates using only the “expected” DAG structure of GO.

As our results indicate, discrepancies in the semantic granularity of gene annotations in knowledgebases represent a significant hurdle to overcome for researchers interested in mining genes based on a set of annotations used in experimental data. As we show, utilizing only the set of specific annotations used in HPA’s experimental data, M2S’s mapping matches only 366 identical gene annotations from the knowledgebase, which comparable to when GOcats is provided with categories matching that set (Figure 4A). GOcats alleviates this problem by allowing researchers to easily define categories at a custom level of granularity so that categories may be specific enough to retain biological significance, but generic enough to encapsulate a larger set of knowledgebase-derived annotations. When we reevaluated the agreement between the raw data and knowledgebase annotations using custom GOcats categories for “cytoskeleton” and “nucleus”, the number of identical gene annotations increased to 776 (Figure 4B).

As GO continues to grow, automated methods to evaluate the structural organization of data will become necessary for curation and quality control. For instance, we recently collaborated in the evaluation of a method to automate auditing of potential subtype inconsistencies among terms in GO [26]. Because GOcats allows versatile interpretation of the GO DAG structure, it has many potential curation and quality control uses, especially for evaluating the high-level ontological organization of GO terms. For example, GOcats can facilitate the integrity checking of annotations that are added to public repositories by streamlining the process of extracting categories of annotations from knowledgebases and comparing them to the original annotations in the raw data. Interestingly, about one-third of the genes annotated with high-confidence in the HPA raw data were missing altogether from the EMBL-EBI knowledgebase when filtered to the HPA-sourced annotations. While this surprised us, the reason appears to be due to HPA’s use of two separate criteria for “supportive” annotation reliability scores and for knowledge-based annotations. For “supportive” reliability, one of several conditions must be met: i) two independent antibodies yielding similar or partly similar staining patterns, ii) two independent antibodies yielding dissimilar staining patterns, both supported by experimental gene/protein characterization data, iii) one antibody yielding a staining pattern supported by experimental gene/protein characterization data, iv) one antibody yielding a staining pattern with no available experimental gene/protein characterization data, but supported by other assay within the HPA, and v) one or more independent antibodies yielding staining patterns not consistent with experimental gene/protein characterization data, but supported by siRNA assay [10,27]. Meanwhile knowledge-based annotations are dependent on the number of cell lines annotated; specifically, the documentation states, “Knowledge-based annotation of subcellular location aims to provide an interpretation of the subcellular localization of a specific protein in at least three human cell lines. The conflation of immunofluorescence data from two or more antibody sources directed towards the same protein and a review of available protein/gene characterization data, allows for a knowledge-based interpretation of the subcellular location” [10,28]. Unfortunately, we were unable to explore these differences further, since the experimental data-based subcellular localization annotations appeared aggregated across multiple cell lines, without specifying which cell lines were positive for each location. Meanwhile, tissue-and cell-line specific data, which contained expression level information, did not also contain subcellular localizations. Therefore, we would suggest that HPA and other major experimental data repositories always provide a specific annotation reliability category in their distilled experimental datasets that matches the criteria used for deposition of derived annotations in the knowledgebases. Such information will be invaluable for performing knowledgebase-level evaluation of large curated sets of annotations. One step better would involve providing a complete experimental and support data audit trail for each derived annotation curated for a knowledgebase, but this may be prohibitively difficult and time-consuming to do.

## Materials and Methods

Materials and methods as well as additional methods illustrations and flowcharts are provided in Supplementary Data 1, 3, 4, 6, 7, 8, 12, and 13.

## Availability and Future Directions

The Python software package GOcats is an open-source project under the BSD-3 License and available from the GitHub repository https://github.com/MoseleyBioinformaticsLab/GOcats. Documentation is available at http://gocats.readthedocs.io/en/latest/. All figures and supplementary data are available on the FigShare repository: https://figshare.com/s/21defede0cd3865742d4 along with the specific version of GOCats used to generate these results: https://figshare.com/s/134d06901a7a6f331934.

We are actively developing the codebase and appreciate any contributions and feedback provided by the community. We are extending the API and adding additional capabilities to handle more advanced annotation enrichment analysis use-cases.

## Acknowledgements

This work was supported in part by grants NSF 1252893 (Moseley), NIH 1U24DK097215-01A1 (Higashi, Fan, Lane, Moseley), and NIH UL1TR001998-01 (Kern). We thank Dr. Robert M. Flight for his advice and expertise regarding the statistics reported in this project, for the generation of the plots in Figure 3, and for his feedback during the drafting of the manuscript. We thank Dr. Thilakam Murali for extensive feedback on the general scientific readability of the manuscript.

## Author Contributions

E.W.H. worked on the design of GOcats, implemented GOcats, performed all analyses, interpreted results, and wrote over half of the manuscript. H.N.B.M. worked on the design of GOcats, troubleshooted various aspects of GOcats implementation, designed the analyses, interpreted results, and wrote a significant portion of the manuscript.

